# A low resolution epistasis mapping approach to identify chromosome arm interactions in allohexaploid wheat

**DOI:** 10.1101/377713

**Authors:** Nicholas Santantonio, Jean-Luc Jannink, Mark E. Sorrells

## Abstract

Epistasis is an important contributor to genetic variance, even in inbred populations where it is present as additive by additive interactions. Testing for epistasis presents a multiple testing problem as the search space for modest numbers of markers is large. Additionally, single markers do not necessarily track functional units of interacting chromatin as well as haplotype based methods do. To harness the power of multiple markers while drastically minimizing the number of tests conducted, we present a low resolution test for epistatic interactions across whole chromosome arms. Two additive genetic covariance matrices are constructed from markers on two different chromosome arms. The Hadamard product of these additive covariance matrices is then used to produce the additive by additive epistasis covariance matrix between the two chromosome arms. The covariance matrices are subsequently used to estimate an epistatic interaction variance parameter in a mixed model framework, while correcting for background additive and epistatic effects. We find significant epistatic interactions for 2% of interactions tested for four agronomic traits in a population of winter wheat. Interactions across homeologous chromosome arms were identified, but were less abundant than other interaction chromosome arm pairs. Of these, homeologous chromosome arm pair 4BL and 4DL showed a strong relationship between the product of their additive effects and the interaction effect that may be indicative of functional redundancy. Several chromosome arms were involved in many interactions across the genome, suggesting that they may contain important large effect regulatory factors. The differential patterns of epistasis across different traits suggests that detection of epistatic interactions is robust when correcting for background additive and epistatic effects in the population. The low resolution epistasis mapping method presented here identifies important epistatic interactions with a limited number of statistical tests at the cost of relatively lower precision.

## 2 Introduction

Epistasis is the interaction of alleles, or variants, at two or more loci. Early observations of epistasis by William Bateson (2007) were mostly qualitative, noting that certain loci could mask the effects at other loci. Quantitative epistasis was first suggested and defined by Ronald Fisher (1919) who coined the term ‘epistasy’. Statistically, epistasis is the deviation from an additive expectation of two or more loci, often described as a change in the slope of one locus based on the genotype at another locus (Fisher 1919). Variance due to quantitative epistasis has been shown to be an important contributor to the genetic variance in populations of model organisms such as *Arabidopsis* (Malmberg et al. 2005; Kusterer et al. 2007), as well as crop species such as maize (Stuber and Moll 1971; Melchinger, Geiger, and Schnell 1986; Lamkey, Schnicker, and Melchinger 1995; Wolf and Hallauer 1997; Lukens and Doebley 1999) and rice (Yu et al. 1997; Li et al. 2008; Shen et al. 2014). Significant epistasis has been reported in allopolyploid crops like cotton (Lee, Cockerham, and Smith 1968) and wheat (Crossa et al. 2010; Jiang et al. 2017). Epistasis across subgenomes may be indicative of interactions between homeologous loci, analogous to dominance in diploids, and a possible contributor to that adaptation of these crops to a wide landscape (Wendel 2000; Adams and Wendel 2005; Chen 2010; Chen 2013). However, there is still little direct evidence that epistasis between homeologous loci is a large contributor to the total genetic variance in allopolyploids.

Epistasis has also been shown to be an important contributor to evolution (Doebley, Stec, and Gustus 1995; Lukens and Doebley 1999; Carlborg et al. 2006; Phillips 2008; Hansen 2013; Doust et al. 2014). There has been considerable effort over the past several decades to incorporate these non-additive genetic factors into the genotype to phenotype map. More recently these effects have been incorporated into whole genome prediction models (Vitezica, Varona, and Legarra 2013; Martini et al. 2016; Jiang and Reif 2015; Huang and Mackay 2016; Akdemir and Jannink 2015; Wolfe et al. 2016; Akdemir, Jannink, and Isidro-Sánchez 2017; Jiang et al. 2017).

In practice, detecting epistatic interactions is difficult. The pairwise search space is large even for modest numbers of markers. For example, a population genotyped with 100 markers would require 4,950 tests for pairwise epistasis. With advances in genotyping technologies, the number of DNA markers available is typically much larger, in the tens to hundreds of thousands, and more recently in the millions. In this study, 11,604 markers were available, which would result in approximately 67 million tests for pairwise epistasis. A 0.05 genome-wide Bonferroni significance threshold for all pairwise epistasis tests in this study would then be 7.4 × 10^−10^.

Several methods have been proposed to reduce the multiple testing problem. Epistasis is partitioned in part to the additive variance, particularly when allele frequencies differ from 0.5 at either locus (Hill, Goddard, and Visscher 2008). Therefore, genome-wide scans can be used to first identify marker alleles (or variants) with a significant additive effect, then test only all pairwise variants identified in the scan (Carlson et al. 2004). This can greatly reduce the number of epistatic tests performed, while increasing the likelihood that epistasis will be identified. Other methods include relaxing the multiple test correction threshold (Benjamini and Hochberg 1995), or reducing the marker pairs tested based on some criteria such as biological function or other filtering methods (Ritchie 2011; Cowman and Koyutürk 2017; Crawford et al. 2017).

The multiple test correction problem is not the only challenge to identifying epistatic interactions. Allele frequency, linkage disequilibrium and the number of alleles at a given locus can all reduce the efficacy of pairwise marker epistasis detection. Low allele frequencies at either locus reduce the epistatic effect, partitioning it to the additive instead (Hill, Goddard, and Visscher 2008). Less than perfect linkage disequilibrium between the markers and causal mutations also reduces the apparent effect size, limiting detection much as it does for additive effects (Carlson et al. 2004). SNP markers are typically considered bi-allelic, despite the potential for many different alleles in the population. The impact of these factors can be reduced by using multiple linked markers to determine haplotypes. Haplotypes have been shown to be powerful in the detection of additive and interaction effects by accurately tracking larger segments of DNA in high or perfect LD, and allowing multiple alleles at every locus (Lin and Zeng 2006; Zhang et al. 2012; Jiang, Schmidt, and Reif 2018). While allele frequencies are typically reduced using haplotypes (i.e. the frequency of two alleles will be higher than the frequency of three alleles), the added power from accurately tracking relevant LD blocks make these methods attractive.

Haplotypes do not need to be assigned directly to gain an advantage from using multiple markers to identify regions associated with complex traits. Regional heritability mapping (Nagamine et al. 2012; Riggio and Pong-Wong 2014) has been used to identify additive effects of rare and common variants in humans (Nagamine et al. 2012; Shirali et al. 2016) as well as plants species like eucalyptus (Resende et al. 2017) and cassava (Okeke et al. 2018). These methods employ the estimation of additive covariance between individuals based on markers in a given region of chromatin, and are used in a mixed model to estimate the genetic variance attributable to the region. Variance components can then be tested to determine if they are greater than zero using a likelihood ratio test.

We propose a method to greatly reduce the number of statistical tests while taking advantage of multiple markers to determine importance of epistatic interactions across chromosome arms of an allohexaploid wheat population. This method is similar to the “divide and conquer” method of Akdemir and Jannink (2015), but models interactions across chromosomes instead of local epistasis. Epistatic covariances can be formed using the Hadamard product of component additive or dominance covariance matrices (Henderson 1985; Jiang and Reif 2015; Martini et al. 2016). Additive by additive epistatic interactions between disjoint sets of related (i.e. linked) markers can be modeled by first calculating an additive covariance for each marker set, *K_i_* and *K_i′_*, and using *K_i_* ⊙ *K_i′_* as the covariance estimate of the epistatic term between these sets. We define marker sets by the chromosome arm to which they belong, and estimate the epistatic variance component between the two arms using Restricted maximum likelihood (REML) while correcting for background additive and epistatic effects.

Common wheat is an important allohexaploid crop with three subgenomes, A, B and D, resulting from hybridization events approximately 500 thousand and 10 thousand years ago. Due to the allopolyploid nature of wheat, we were interested in identifying interactions across homeologous loci. Interactions at homeologous loci are analogous to dominance effects in diploid hybrids, and could be used to fix favorable homeoallelic interactions in inbred lines (Wendel 2000; Adams and Wendel 2005; Birchler et al. 2010; Chen 2010; Chen 2013). Of the 21 chromosomes of wheat, chromosome arms pairs include 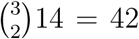 homeologous pairs, 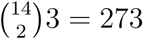 within subgenome pairs, and 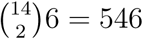 across subgenome arm pairs.

Each chromosome arm of the wheat genome was sequenced independently using flow cytometry to assist in the assembly of the large complex genome (International Wheat Genome Sequencing Consortium 2014). The lone exception was chromosome 3B, which was sequenced and assembled in its entirety before the other chromosomes of wheat (Paux et al. 2008; Choulet et al. 2014). Therefore, assigning markers to a chromosome arm is feasible, but their position along that arm may not be well defined if the number of scaffolds is large, as was the case with the first wheat survey sequence (International Wheat Genome Sequencing Consortium 2014). Using markers across an entire chromosome arm known to be homeologous to another chromosome arms may therefore be a better strategy than attempting to assign single homeologous marker pairs. If interactions are detected across homeologous regions, this may provide evidence of beneficial homeoallelic interactions indicative of inter-genomic heterosis.

We demonstrate the low resolution epistasis mapping methodology using the CNLM winter wheat dataset from Santantonio et al. (2018a; 2018b), and show that epistasis can be detected between homeologous and non-homeologous chromosome arms.

## 3 Materials and Methods

### 3.1 Chromosome centromere positions

Chromosome centromere positions were provided by the IWGSC for all chromosomes except 3B (IWGSC, personal communication, March 1, 2017). Those positions were assigned by determining where chromosome arm library reads aligned to the final assembly. Each chromosome arm was sequenced independently using flow-cytometry to remove the chromosome arm from a series of aneuploid stocks, each containing an extra arm. The lone exception was chromosome 3B, which was sequenced in its entirety, so no centromere position was available for the 3B chromosome. Centromere start and stop addresses provided by IWGSC are shown in Supplemental Table S1.

While restriction sites are expected to be uniformly distributed throughout the genome, methylation of cytosine is not. One of the restriction enzyme used to generate GBS libraries, MspI, is sensitive to DNA methylation, digesting unmethylated DNA at a much higher rate than methylated DNA (McClelland, Nelson, and Raschke 1994). Methylation is an important regulator of chromatin structure, where euchromatin tends to contain few methylation sites relative to heterochromatin (Keshet, Lieman-Hurwitz, and Cedar 1986). Therefore restriction sites in heterochromatin with high levels of methylation, such as at the centromere, are less likely to be retained as GBS markers because digestion is less likely to happen at these sites. This means that the GBS markers can be used to roughly assign a centromere position using the density of GBS markers along the chromosome.

To determine the centromere position of 3B, we employed kernel density estimation using the `density()` function of the ‘stats’ package in R to determine the smoothed density of GBS marker positions. We then assigned the 3B centromere interval to the chromosome positions flanking the second position for which the derivative of the density was zero. We performed this operation for all chromosomes to determine the efficacy of this method for determining the centromere position.

### 3.2 Chromosome arm resolution epistasis

The low resolution epistasis mapping approach employed here uses markers from two defined regions, *i* and *i*′, to calculate additive covariance between individuals based on those regions (i.e **K**_*i*_ and **K**_*i*′_ ∀ *i* ≠ *i*′). The Hadamard product of these additive covariance matrices can be used to produce the pairwise additive by additive epistatic relationship, **K**_*i*×*i*′_ = **K**_*i*_ ⊙ **K**_*i*′_, between these two regions (Henderson 1985; Martini et al. 2016). In this study, we defined regions as the short (*S*) and long (*L*) arms of each chromosome, where *i* ϵ {1*AS*, 1*AL*, 1*BS*, …, 7*BL*, 7*DS*, 7*DL*}. Variance components for each region and their respective interaction were estimated by fitting the following nested models

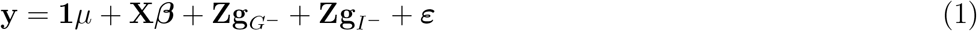

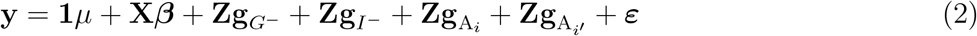

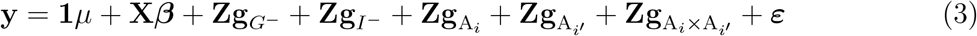

where 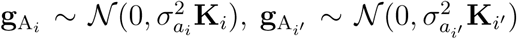 and 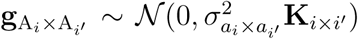. *g_G^−^_* and *g_I^−^_* were modeled as previously described in Santantonio et al. (Santantonio, Jannink, and Sorrells 2018b, equation 5), but with markers belonging to region *i* and *i′* removed prior to calculating the covariances.

Sequential nested likelihood ratio tests were used to determine if the additive (model 2 versus model 1) and interaction (model 3 versus model 2) variance estimates of the chromosome arms were greater than zero. From the Neyman-Pearson lemma (Neyman and Pearson 1933), the likelihood ratio test statistic is defined as 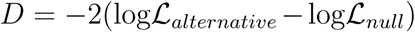, where 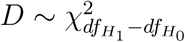, and is uniformly most powerful (UMP).

BLUPs were subsequently used to look for patterns between additive and interaction effects for the chromosome arm pair. The pairwise product of the additive chromosome arm BLUPs was then compared to the chromosome arm interaction BLUPs, in a manner analogous to the Additive × Additive single locus model (Hill, Goddard, and Visscher 2008; Santantonio, Jannink, and Sorrells 2018a). Negative associations should indicate a less than additive model, whereas positive relationships would demonstrate a more than additive epistatic effect.

When a variance parameter is very close to zero, parameter estimates are unreliable, and were therefore considered to be zero. Markers were oriented by minor allele frequency as it was unclear how to apply the previous orientation schemes to multiple sets of markers. It is not clear what effect, if any, this orientation had on the estimates of interaction effects.

For the 14 three-way homeologous arm sets, a three-way interaction was included and tested against a model with only the three two-way interaction terms. We did not attempt to run all three-way chromosome arm combinations, as this would have been computationally infeasible, with 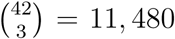 combinations. The Hadamard product of the three additive covariance matrices was used to produce the three-way additive by additive by additive epistatic relationship, **K**_*i*×*i′*×*i″*_ = **K**_*i*_ ⊙ **K**_*i′*_ ⊙ **K**_*i″*_. The following two models were fit to test the three-way interaction.

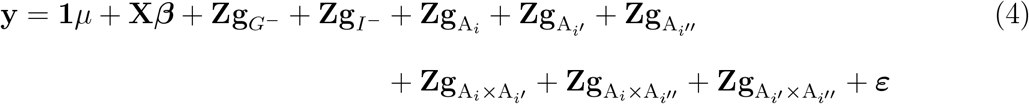

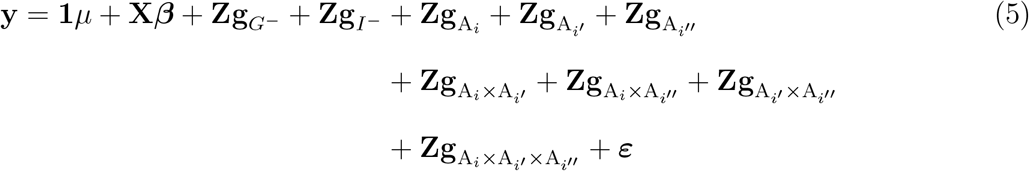

The likelihood ratio test was then used to determine if adding the three-way interaction term significantly improved the model fit beyond the two-way interaction terms.

### 3.3 Software

Variance component estimation was accomplished using Restricted Maximum Likelihood (REML) inmplemented in ASReml-R (Gilmour 1997; Butler 2009). Other computation, analyses and figures were made using base R (R Core Team 2015) implemented in the Microsoft Open R environment 3.3.2 (Microsoft 2017) unless noted otherwise. The ‘circlize’ 204 R package (Gu et al. 2014) were used to make Figures S3 and 3. LATEXtables were generated using The R package ‘xtable’ (Dahl 2016).

### 3.4 Data Availability

All data used for this study can be found in (Santantonio, Jannink, and Sorrells 2018b).

## 4 Results

### 4.1 Centromere positions

Most of the GBS marker density estimates of centromere locations agreed well with the positions provided by the IWGSC (Supplemental Figure S1). Chromosomes 1D and 4A were exceptions. We estimated the 3B centromere to be positioned between 347.3 Mbp and 347.9 Mbp (Supplemental Table S1).

### 4.2 Model fit and p-value distribution

Homeologous chromosome arm pair models each had five random genetic effects and therefore five covariance structures for the two-way interaction models. All models converged, but some variance parameter estimates were often close to the parameter boundary and were considered to be zero. Variance component estimates on the boundary did not occur for the background additive or epistatic effects, but often occurred for one or both of the additive chromosome arm effects or the interaction effect. This resulted in a relatively large number of additive and interaction variance component tests with a p-value of 1. As a result, p-value distributions were heavily skewed toward 0 and 1 (Figure S2 and Supplemental Figure 1). Most chromosome arms had low additive effect p-values, whereas most interaction p-values were high, indicating that the majority of chromosome arm pairs do not have effect interactions large enough to detect.

**Figure 1:**
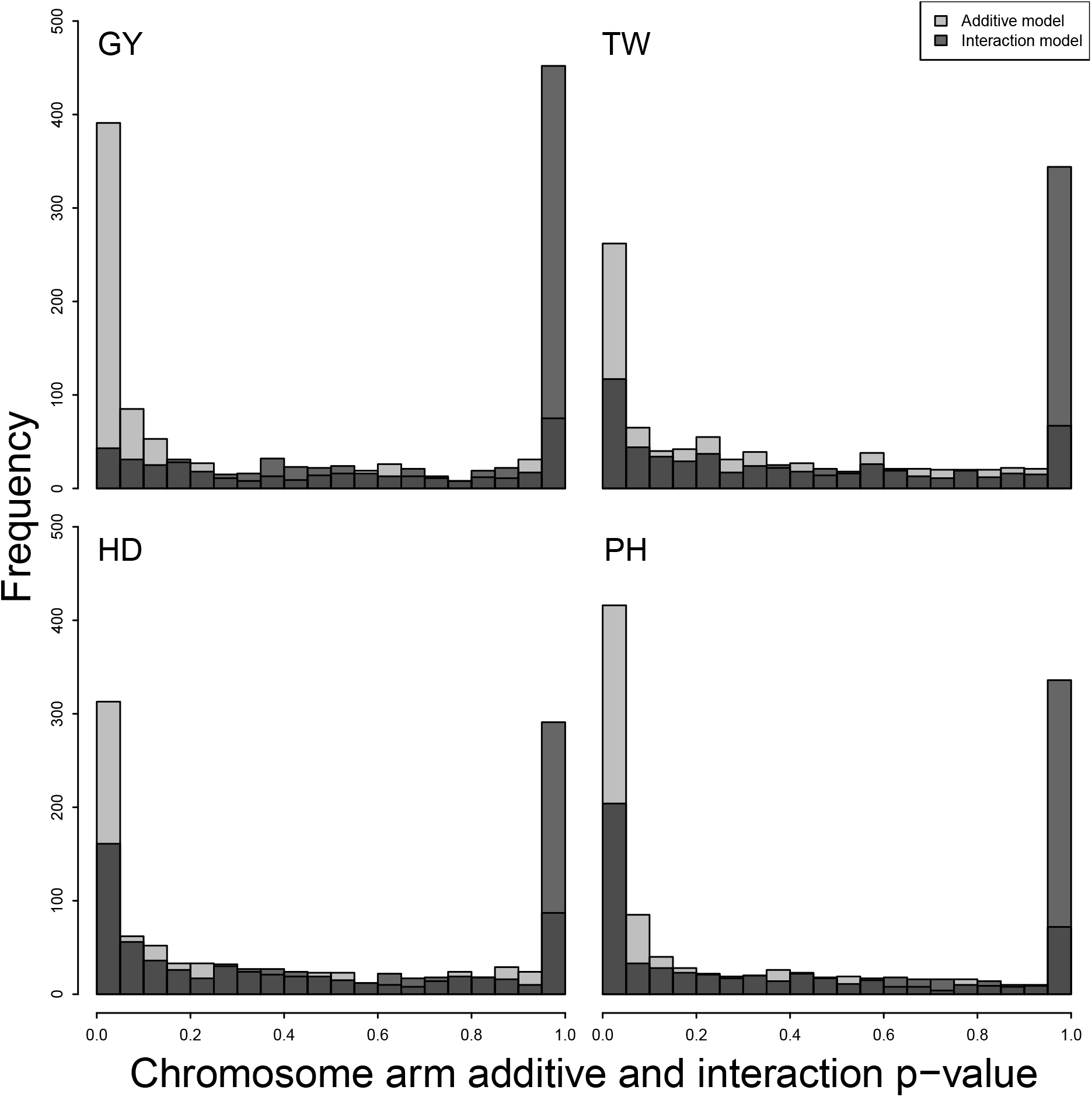
Distribution of p-values for all 861 possible chromosome arm pair models for four traits, GY, TW, PH and HD. The p-value from the likelihood ratio test for the additive chromosome arm model is plotted in light gray, whereas the p-value from the the interaction model test is shown in dark gray.

### 4.3 Homeologous arm tests

**Table 1:** Table of significant homeologous chromosome arm interactions. The proportion of genetic variance attributed to each arm and their corresponding interaction are shown with statistical significance from a nested likelihood ratio test.

| Trait | arm<sub>i</sub> | arm<sub>i'</sub> | ($h^2_{\text{arm}_i}, h^2_{\text{arm}_{i'}}$)<sup>a</sup> | $h^2_{\text{arm}_i \times \text{arm}_{i'}}$ | $\rho^b$ |
| --- | --- | --- | --- | --- | --- |
| GY | 5BS | 5DS | (0.038, 0) | 0.028** | 0.27*** |
| GY | 7AL | 7BL | (0.018, 0) | 0.041* | 0.1*** |
| PH | 2AS | 2DS | (0.021, 0.079)*** | 0.033*** | -0.04 |
| PH | 4AS | 4DS | (0, 0.039)*** | 0.017* | 0.19*** |
| PH | 4AL | 4BL | (0.013, 0.034)* | 0.029* | 0.1*** |
| PH | 4AL | 4DL | (0.015, 0.0038) | 0.027*** | 0.07** |
| PH | 4BS | 4DS | (0.0021, 0.031)*** | 0.049*** | 0.18*** |
| PH | 4BL | 4DL | (0.048, 0.0033)* | 0.058*** | -0.65*** |
| PH | 6AL | 6DL | (0.11, 0.0053)** | 0.024* | 0.06* |
| PH | 7AL | 7BL | (0, 0.07) | 0.029** | 0.45*** |
| TW | 1BS | 1DS | (0, 0) | 0.073*** | 0 |
| TW | 4BL | 4DL | (0.096, 0.049)*** | 0.013* | 0.14*** |
| TW | 6AL | 6BL | (0.031, 0) | 0.047* | 0.12*** |
| TW | 7AL | 7DL | (0.019, 0.03) | 0.14*** | -0.04 |
| TW | 7BL | 7DL | (0.043, 0.061) | 0.092*** | 0.16*** |
| HD | 1BS | 1DS | (0, 0) | 0.018* | 0 |
| HD | 4BS | 4DS | (0, 0.0023) | 0.014** | 0.15*** |
| HD | 6AS | 6BS | (0.0078, 0.041)* | 0.049*** | 0.02 |
| HD | 6AS | 6DS | (0.014, 0) | 0.046*** | -0.03 |
| HD | 6AL | 6BL | (0.0087, 0.11)* | 0.013* | -0.21*** |
| HD | 7AS | 7DS | (0.013, 0.045)** | 0.032* | -0.05* |
| HD | 7AL | 7BL | (0, 0.045)*** | 0.025* | 0.14*** |
| HD | 7BS | 7DS | (0.013, 0.054)*** | 0.012* | 0.29*** |

<sup>a</sup> $h^2$ represents the proportion of the chromosome arm additive or interaction variance component estimates to the total genetic variance.
<sup>b</sup> $\rho$ indicates the correlation between the product of the additive arm effects and their interaction effect with correlation coefficients significantly different from zero indicated by stars. If only one additive effect had a non-zero variance, the correlation coefficient shown is the correlation between the additive effect with the non-zero variance and the interaction effect.
\*, \*\*, and \*\*\* correspond to p-values < 0.05, 0.01, and a Bonferroni correction of 0.05/42 = 0.0012, respectively.

The U-shaped distribution of the p-values suggested that when the true variance was very small or zero, the average information algorithm estimated the parameter on the boundary (i.e. 0), and when it was positive, the p-value tended to be low. Larger sample sizes may be necessary to obtain uniform p-value distributions when the null hypothesis is true. We therefore considered all homeologous arm pairs with an interaction variance p-value less than 0.05 that also had positive additive variance component estimates to determine the relationship between additive chromosome arm effects and their interaction.

Seventeen homeologous chromosome arm pairs had significant interaction effects for at least one of the four traits (Table 1 and Supplemental Figure S3). Interactions involving homeologs 4 and 7 were overrepresented, with 14 of the 22 significant interactions identified between one of these two homeologs. Despite significant pairwise homeologous marker interactions found on chromosome homeologs 1 and 5 for HD and homeolog 3 for PH (Santantonio, Jannink, and Sorrells 2018a), chromosome arm pair tests failed to detect the significant homeologous marker set interactions on those arms. The failure to detect these significant regions using the chromosome arm test suggests that these are either spurious associations, or their signal is being washed out by the abundance of uninformative markers on those chromosome arms. The lack of a two-way arm interactions PH interaction on chromosome arm 3S agrees with the homeologous marker set identified there, where only the three-way homeologous marker set interaction term was significant.

The test for three-way homeologous chromosome arm interactions only revealed three sets of homeologous arms that had a significant three-way interaction (Table 2). The three-way 3S chromosome arm interaction for PH was found to have a positive three-way arm interaction variance parameter estimate with a p-value of p = 0.02, supporting the evidence from the homeologous marker set on 3S. The 7L three-way arm interaction term was also found to have a low p-value for TW of p = 0.006, also confirming the significant three-way homeologous marker set found there.

Many interactions were detected on chromosome arms where no homeologous marker sets were identified with a significant interaction effect. Notably, a strong interaction effect was identified on homeolog 6S for HD, and two regions for GY on 5S and 7L, where no significant homeologous interaction sets were identified. Neither of the interacting pairs for GY had a p-value lower than a homeologous arm Bonferroni correction of 0.05 / 42 = 0.0012.

Relationships between chromosome arm additive and interaction effects were only considered for the ten chromosome arm pair trait combinations that had all chromosome arm additive and interaction effects with significant non-zero variance components. Of these ten, six had significant correlations between the additive product and the interaction with an absolute value ≥ 0.1. Four of these showed positive relationships, while the other two showed negative relationships. By far the strongest relationship detected was between 4BL and 4DL for PH (*ρ* = −0.65, Figure 2), indicating that individuals with high or low additive values for both arms tended to have genotypic values less than expected by additivity. Conversely, the same 4BL/4DL pair had a weak, yet positive relationship for TW (*ρ* = 0.14, Supplemental Figure 2). The 4BS/4DS pair, where the *Rht-1* genes are known to reside, had a weak, yet significant, positive correlation for PH (Supplemental Figure S5).

### 4.4 All pairwise arm tests

For all 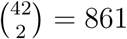 pairwise chromosome arm pairs, we only consider those tests that passed a Bonferroni threshold of 0.05/861 = 5.8 × 10^−5^ in this section. Seventy nine chromosome arm interaction variance components were declared significantly greater than zero for at least one trait, representing about 2% of the number tested (Table 4). Of these, interactions for the PH trait were the most prevalent, representing 49 (62%), of the interactions detected. HD and TW accounted for the remaining 13 (16%) and 17 (22%) interactions. No chromosome arm interactions were detected for GY at the Bonferroni significance threshold. No interactions were detected for any of the traits involving chromosome arms 1AS, 1DL, 2AS, 2DL, 3DL, 4AS, 5AS, 5BL, 5DL, 6BL, 6DS, and 7BS at this threshold.

There were several chromosome arms that appeared to be interacting with multiple loci (Supplemental Table S2). Of these, several clearly stand out (Figure 3). Chromosome arms 1AL, 2AL, 2DS, 4BS, 4DS, 4DL, 6AS and 7AL were involved in five or more interacting pairs for PH, with 2DS, 4DS and 4DL. The 4D chromosome in particular was involved in almost half (21) of the interacting arm pairs for this trait. 7DL was involved in all but three of the interacting pairs detected for the TW trait. Arm interactions for HD did not cluster to one or a few arms in the same way as PH and TW, but 6AS and 7BL were each involved in five interacting pairs for this trait.

**Figure 2:**
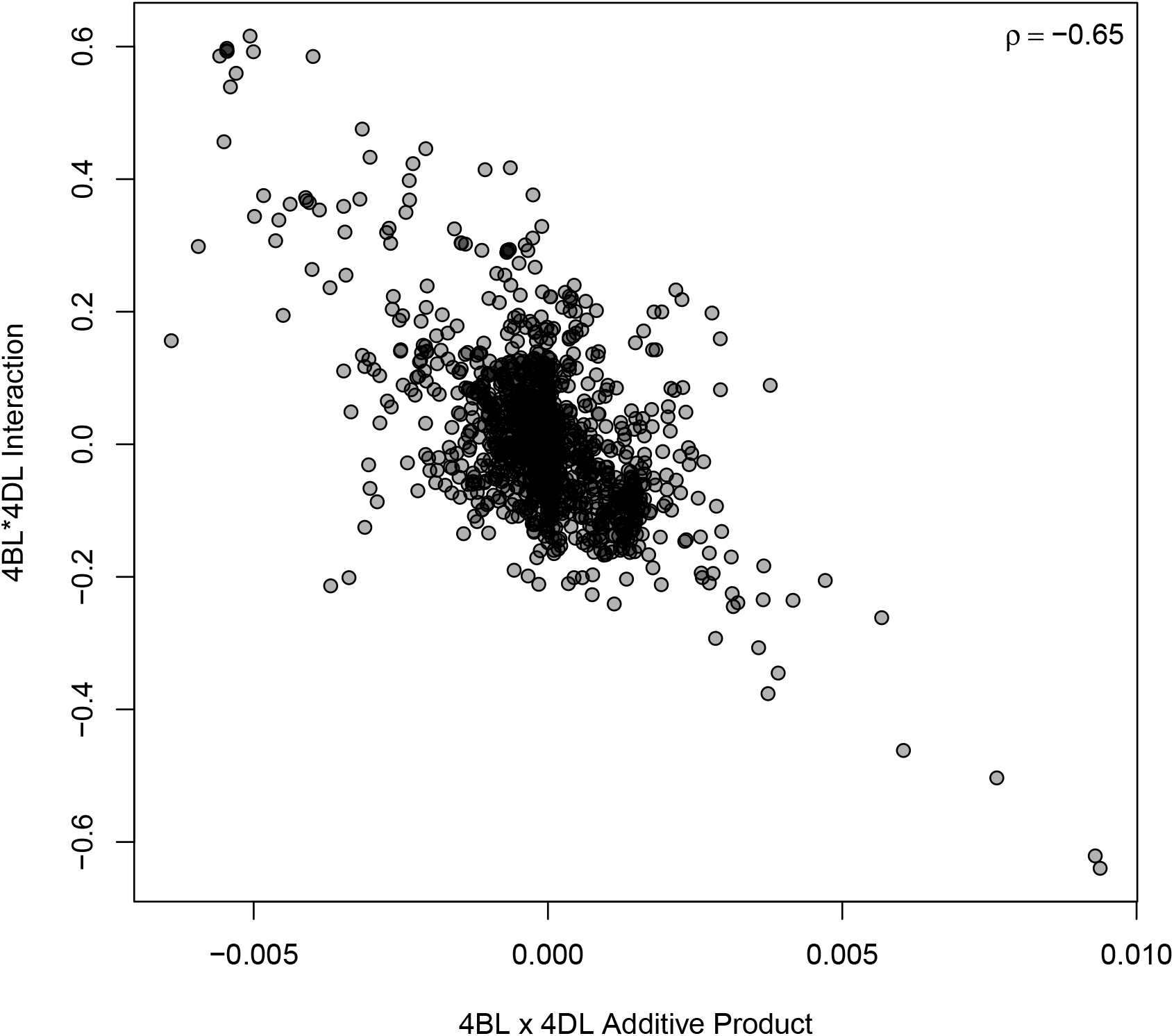
Interaction effect of chromosome 4BL by 4DL plotted against the product of the additive effects for 4BL and 4DL for PH. *ρ* indicates the Pearson correlation coefficient.

**Table 2:**
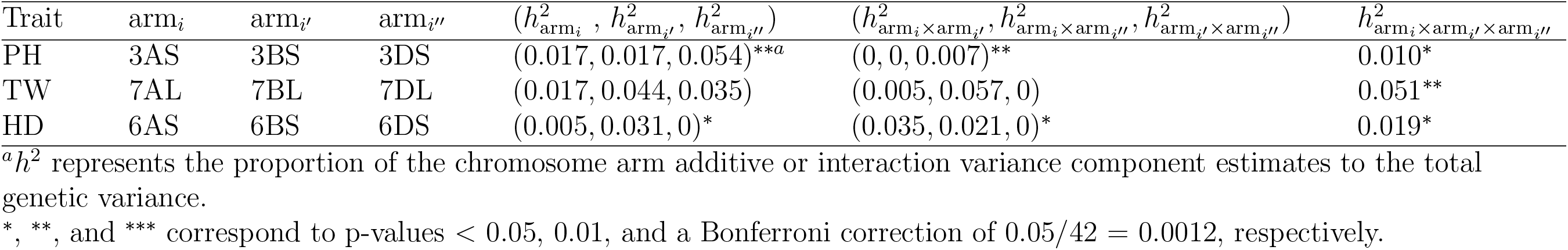
Table of significant three-way homeologous chromosome arm interactions. The proportion of genetic variance attributed to each arm and their corresponding interaction are shown with statistical significance from a nested likelihood ratio test indicated by stars.

**Table 3:** Table of significant chromosome arm interactions for all four traits. The proportion of genetic variance attributed to each arm and their corresponding interaction are shown with statistical significance from a nested likelihood ratio test.

| Trait | arm<sub>i</sub> | arm<sub>i'</sub> | ($h^2_{\text{arm}_i}, h^2_{\text{arm}_{i'}}$)<sup>a</sup> | $h^2_{\text{arm}_i \times \text{arm}_{i'}}$ | $\rho^b$ |
| --- | --- | --- | --- | --- | --- |
| PH | 1AL | 2AL | (0.039, 0.06)****<sup>a</sup> | 0.09**** | 0.37*** |
| PH | 1AL | 3AS | (0.065, 0.011)** | 0.076**** | -0.01 |
| PH | 1AL | 7AL | (0.063, 0)* | 0.067**** | 0.35*** |
| PH | 1AL | 4BS | (0.042, 0.024)** | 0.073**** | 0.08** |
| PH | 1AL | 4BL | (0.066, 0.012)*** | 0.086**** | -0.05 |
| PH | 1AL | 7BL | (0.091, 0.043)*** | 0.059**** | 0.01 |
| PH | 1AL | 2DS | (0.079, 0.094)**** | 0.065**** | 0.18*** |
| PH | 1AL | 4DS | (0.048, 0.037)**** | 0.064**** | -0.04 |
| PH | 1AL | 4DL | (0.07, 0.002)** | 0.048**** | 0.05 |
| PH | 2AL | 7AL | (0.04, 0)** | 0.056**** | 0.27*** |
| PH | 2AL | 4BS | (0.06, 0.032)*** | 0.09**** | -0.17*** |
| PH | 2AL | 2DS | (0.036, 0.097)**** | 0.061**** | -0.17*** |
| PH | 2AL | 4DL | (0.039, 0)*** | 0.069**** | 0.23*** |
| PH | 3AS | 6AS | (0.015, 0.069)** | 0.084**** | -0.22*** |
| PH | 3AS | 6AL | (0, 0.086)** | 0.07**** | 0.35*** |
| PH | 3AS | 7AL | (0.0081, 0) | 0.06**** | 0.27*** |
| PH | 3AS | 7BL | (0.00084, 0.038)* | 0.068**** | 0.01 |
| PH | 3AS | 4DS | (0.02, 0.033)**** | 0.057**** | -0.09** |
| PH | 3AL | 2DS | (0.0042, 0.068)** | 0.042**** | -0.11*** |
| PH | 3AL | 4DS | (0.0075, 0.033)**** | 0.05**** | -0.17*** |
| PH | 3AL | 4DL | (0.0015, 0.0097) | 0.023**** | -0.07** |
| PH | 6AS | 7AL | (0.083, 0)** | 0.069**** | 0.4*** |
| PH | 6AS | 1BS | (0.078, 0.011)*** | 0.1**** | 0.02 |
| PH | 6AS | 4BS | (0.054, 0)*** | 0.13**** | -0.01 |
| PH | 6AS | 4DS | (0.16, 0.042)**** | 0.052**** | 0.03 |
| PH | 6AS | 4DL | (0.087, 0)** | 0.073**** | 0.23*** |
| PH | 6AS | 6DL | (0.069, 0.0077)** | 0.084**** | 0.09** |
| PH | 7AL | 4BL | (0, 0.029)* | 0.058**** | 0.29*** |
| PH | 7AL | 2DS | (0.00064, 0.064)** | 0.054**** | 0.11*** |
| PH | 7AL | 6DL | (0, 0.016) | 0.051**** | 0.31*** |
| PH | 1BS | 2DS | (0, 0.11)*** | 0.062**** | 0.22*** |
| PH | 1BS | 4DS | (0, 0.035)**** | 0.042**** | 0.26*** |
| PH | 1BS | 4DL | (0.023, 0.0091) | 0.041**** | 0.3*** |
| PH | 2BS | 4BS | (0.027, 0.053)* | 0.084**** | -0.11*** |
| PH | 2BS | 4DS | (0.038, 0.032)****<sup>a</sup> | 0.069**** | -0.12*** |
| PH | 2BL | 4DS | (0.2, 0.029)**** | 0.052**** | 0.13*** |
| PH | 4BS | 4DS | (0.0021, 0.031)**** | 0.049**** | 0.18*** |
| PH | 4BS | 4DL | (0, 0) | 0.056**** | 0 |

<sup>a</sup> $h^2$ represents the proportion of the chromosome arm additive or interaction variance component estimates to the total genetic variance.
<sup>b</sup> $\rho$ indicates the correlation between the product of the additive arm effects and their interaction effect with correlation coefficients significantly different from zero indicated by stars. If only one additive effect had a non-zero variance, the correlation coefficient shown is the correlation between the additive effect with the non-zero variance and the interaction effect.
\*, \*\*, and \*\*\* correspond to p-values < 0.05, 0.01, and a Bonferroni correction of 0.05/42 = 0.0012, respectively.

**Table 4:** Continuation of Table 3 of significant chromosome arm interactions.

| Trait | arm<sub>i</sub> | arm<sub>i'</sub> | ($h^2_{\text{arm}_i}, h^2_{\text{arm}_{i'}}$)<sup>a</sup> | $h^2_{\text{arm}_i \times \text{arm}_{i'}}$ | $\rho^b$ |
| --- | --- | --- | --- | --- | --- |
| PH | 4BS | 6DL | (0, 0.021) | 0.11**** | 0.07** |
| PH | 4BL | 4DS | (0.063, 0.029)**** | 0.051**** | -0.52*** |
| PH | 4BL | 4DL | (0.048, 0.0033)* | 0.058**** | -0.65*** |
| PH | 7BL | 2DS | (0.03, 0.09)*** | 0.071**** | 0 |
| PH | 7BL | 4DL | (0.031, 0.00081)* | 0.094**** | -0.12*** |
| PH | 2DS | 3DS | (0.058, 0.047)**** | 0.034**** | 0.07** |
| PH | 2DS | 4DS | (0.13, 0.038)**** | 0.046**** | 0.07** |
| PH | 2DS | 4DL | (0.11, 0.0057)*** | 0.031**** | 0.04 |
| PH | 2DS | 5DS | (0.08, 0.0031)** | 0.031**** | 0.11*** |
| PH | 4DS | 6DL | (0.031, 0.0023)**** | 0.036**** | 0.08** |
| PH | 4DL | 6DL | (0.0018, 0.011) | 0.026**** | -0.38*** |
| TW | 1AL | 1BS | (0.015, 0) | 0.13**** | 0.23*** |
| TW | 1AL | 7DL | (0.015, 0.061) | 0.097**** | -0.13*** |
| TW | 2AL | 3BS | (0.013, 0) | 0.13**** | 0.19*** |
| TW | 2AL | 7DL | (0.032, 0.063) | 0.11**** | 0 |
| TW | 3AL | 7DL | (0.019, 0.072) | 0.085**** | 0.01 |
| TW | 4AL | 7DL | (0.054, 0.053)* | 0.11**** | -0.11*** |
| TW | 5AL | 1BS | (0.031, 0) | 0.15**** | 0.26*** |
| TW | 5AL | 7DL | (0.026, 0.026)* | 0.14**** | 0.06* |
| TW | 6AS | 7DL | (0, 0.058) | 0.11**** | 0.04 |
| TW | 7AS | 7DL | (0.007, 0.059) | 0.09**** | 0.01 |
| TW | 7AL | 7DL | (0.019, 0.03) | 0.14**** | -0.04 |
| TW | 1BS | 7DL | (0, 0.018) | 0.18**** | 0.18*** |
| TW | 2BL | 7DL | (0.0042, 0.063) | 0.11**** | -0.21*** |
| TW | 3BS | 7DL | (0, 0.071) | 0.12**** | 0.11*** |
| TW | 3BL | 7DL | (0.0026, 0.054) | 0.16**** | 0.11*** |
| TW | 6BS | 7DL | (0, 0.056) | 0.093**** | 0.08** |
| TW | 7DS | 7DL | (0, 0.057) | 0.088**** | -0.02 |
| HD | 6AS | 7AS | (0.0042, 0.008) | 0.098**** | -0.03 |
| HD | 6AS | 1BS | (0.007, 0) | 0.058**** | 0.11*** |
| HD | 6AS | 3BS | (0.012, 0.064)*** | 0.082**** | -0.12*** |
| HD | 6AS | 5BS | (0.0036, 0.00018) | 0.051**** | 0.11*** |
| HD | 6AS | 7BL | (0.0076, 0.033)*** | 0.075**** | 0.06* |
| HD | 7AS | 7BL | (0.0089, 0.037)**** | 0.073**** | -0.05 |
| HD | 7AS | 1DS | (0.013, 0) | 0.053**** | -0.09** |
| HD | 1BS | 4DS | (0, 0.0019) | 0.044**** | 0 |
| HD | 1BL | 4DS | (0, 0.0038) | 0.035**** | 0.02 |
| HD | 7BL | 3DS | (0.036, 0.0015)*** | 0.061**** | 0.03 |
| HD | 7BL | 4DS | (0.031, 0.0028)*** | 0.053**** | 0.05* |
| HD | 7BL | 6DL | (0.035, 0.0098)*** | 0.056**** | -0.15*** |
| HD | 3DS | 4DS | (0, 0.003) | 0.029**** | 0 |

<sup>a</sup> $h^2$ represents the proportion of the chromosome arm additive or interaction variance component estimates to the total genetic variance.
<sup>b</sup> $\rho$ indicates the correlation between the product of the additive arm effects and their interaction effect with correlation coefficients significantly different from zero indicated by stars. If only one additive effect had a non-zero variance, the correlation coefficient shown is the correlation between the additive effect with the non-zero variance and the interaction effect.
\*, \*\*, and \*\*\* correspond to p-values < 0.05, 0.01, and a Bonferroni correction of 0.05/42 = 0.0012, respectively.

Most correlations between the additive products and the epistatic effect were low in magnitude (i.e. < 0.3), particularly for the TW and HD traits. Notable exceptions include the 4BL/4DL pair for PH, which had a highly negative correlation, as previously noted. Pairs with moderate magnitude tended to also include the 4DL chromosome, but other pairs with moderate correlations between the product of their additive and interaction effects included the 1AL/2AL, 1AL/7AL, and 3AS/6AS arm pairs.

## 5 Discussion

### 5.1 Centromere positions

While my assigned position for the 3B centromere position is an estimate, most of the chromosome estimates on other chromosomes were close to the known centromere position. The centromere position estimate reported here should be sufficient to assign most of the 3B markers to the correct chromosome arm for the subsequent analyses.

### 5.2 Model fit and p-value distribution

The distribution of p-values from the likelihood ratio test should be uniform if no true interactions exist. If interactions are important, then we would expect to see a skewed distribution with many small p-values. However, the p-values were often calculated to be one because the variance components were estimated on the parameter boundary (i.e. zero), resulting in the U-shaped distribution. When variance parameters are estimated on the parameter boundary, the p-value becomes one simply due to the fact that the variance component is zero. This is likely due to a lack of sufficient population size to distinguish and resolve multiple small variance components. Perhaps another explanation may be provided by the use of the the average information algorithm to fit the mixed model, which may lose a small portion of information by avoiding the calculation of the second derivative of the likelihood function. While other algorithms exist for solving REML problems, the computational burden of resolving multiple variance components with dense covariance structures may be restrictive. Further investigation is necessary to determine how large a population need be to resolve multiple genetic variance parameters with magnitudes of 1% or less of the total variance.

**Figure 3:**
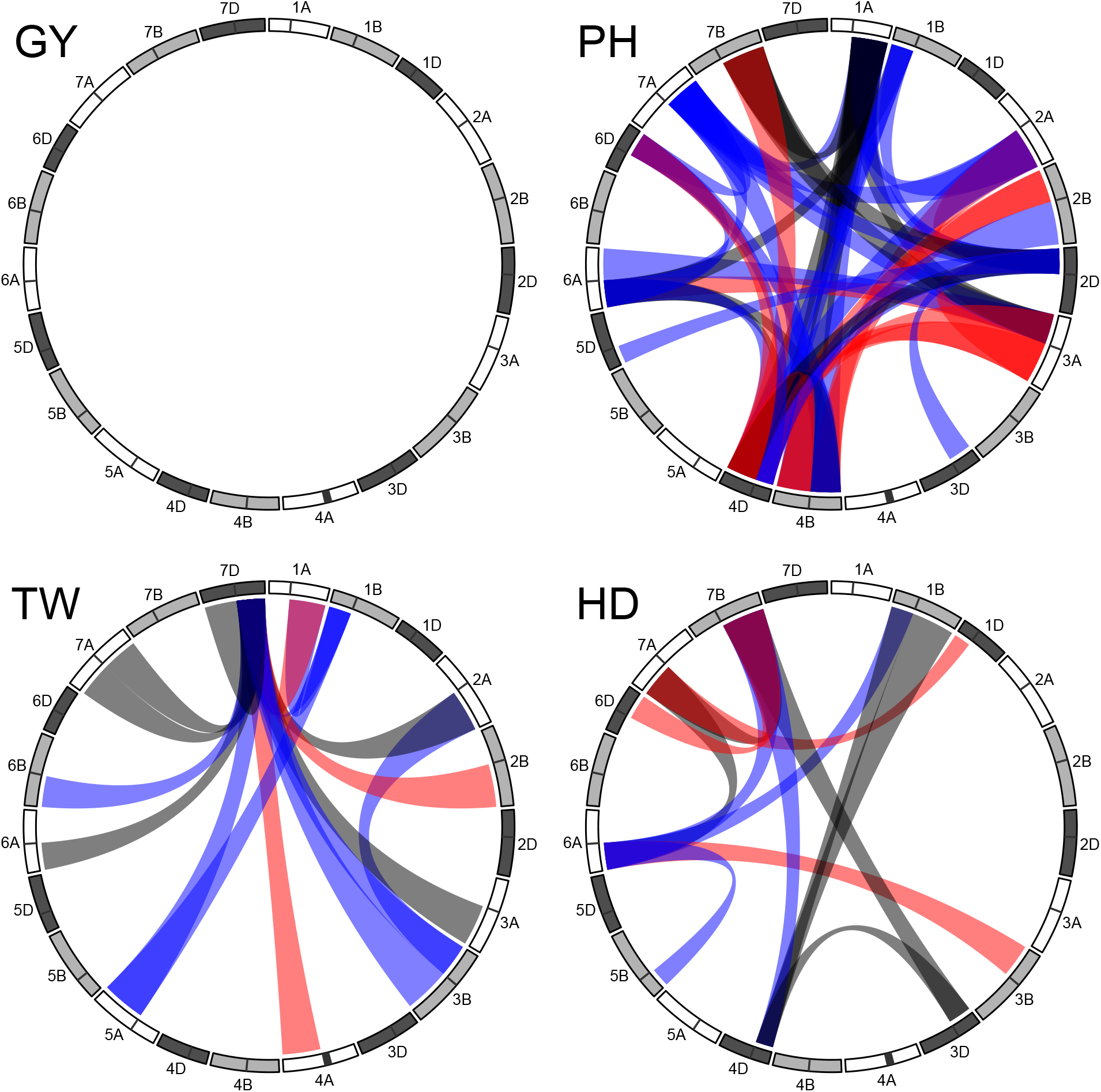
Chromosome arm interactions significant at a Bonferroni correction of 0.05/861 = 5.8 × 10^−5^. Blue and red bridges indicate interactions with a significant positive or negative correlation between the product of the additive effects and their interaction effect, respectively. Black bridges indicate significant interactions that did not have a significant correlation between additive products and the interaction effect.

### 5.3 Homeologous arm tests and additive interaction effect relationships

Most of the homeologous chromosome arm interactions detected across all traits involved homeologs 4 and 7. The less than additive trend observed for the 4BL/4DL pair for PH may suggest a significant degree of gene functional redundancy between these two arms. Despite having a weak positive additive genetic trait correlation between PH and TW (Santantonio, Jannink, and Sorrells 2018b, p. 0.3), the 4BL/4DL pair had a weak, yet positive relationship for TW. This provides evidence that the pattern is not simply a genetic artifact and may indicate differential gene function for these two traits.

A negative correlation for PH was not observed for the 4BS/4DS chromosome arm pair, as might be expected from previous results for the *Rht-1* genes that reside on those chromosome arms. This casts some doubt on the usefulness of these correlations to infer the direction of the epistatic effect. It is unclear if the missing double-dwarf genotype is contributing to this positive correlation in the CNLM population. The relationship between the product of the additive effects and the interaction was thought to mirror the {–1, 1} Additive × Additive epistatic model using a multi-locus approach, but it is unclear what is driving these trends.

For inbred allopolyploids, multi-subunit protein complexes can be comprised of genes from a single subgenome, or from multiple subgenomes. If functional copies of subunits exist on both genomes, the formation of subgenome hetero-complexes may occur. Protein complexes comprised of evolutionarily divergent subunits may have increased or, more likely, decreased functionality. If heterocomplexes display decreased functionality, then we would expect the relationship between the additive and epistatic effects to be negative.

It is unlikely that all homeologous interactions are so large in effect that they are quickly fixed after the hybridization event. The distribution of epistatic effects is likely similar in shape to the distribution of additive effects. These distributions will change based on the complexity of the trait. If a trait is governed by relatively few loci, the relatively few epistatic interactions could have larger effects, and may be easier to detect. In contrast, a distribution of effects with many small non-zero effects may have many more non-zero epistatic effects, but are too small to detect.

### 5.4 All pairwise arm tests

PH appears to exhibit a higher degree of epistasis than either TW or HD. However, the number of interacting loci or chromosome arms detected was not directly related to the observed increase in genomic prediction accuracy (Santantonio, Jannink, and Sorrells 2018b). HD had the largest percent increase in accuracy from the additive only model by including all pairwise additive by additive interactions, yet had the fewest detectable interacting chromosome arms, other than GY. GY showed no evidence of epistasis controlling the trait. This may be due to one of two explanations. The first and most obvious is that grain yield is not subject to epistatic gene action. This would mean that all genes contribute additively to the collection and allocation of resources to vegetative tissue, and then reallocation to the ear during flowering and grain fill. The second and more likely explanation is that GY is the culmination of essentially all the genes working in concert to produce the final outcome, and interactions with such small effects may simply be too small to detect (Xu and Jia 2007; Wu, Chang, and Jing 2012).

While we corrected for population structure on both the additive and epistatic levels (i.e using additive and additive by additive genetic covariance terms), it is possible that residual structure is causing these observed relationships. The drastically different patterns in the arm pair test results for each trait suggests otherwise. If these interactions were due to population structure, we would expect to see similar patterns of significance across all traits. When we omitted the background epistatic effect, most of the 861 interactions were significant (results not shown). We deemed this to be due to chromosome arm epistatic relationship matrices modeling close relationships in the population regardless of which unit of chromatin was used to determine those relationships. However, it is possible that these interactions are far more prevalent than suggested here, and that correction for background epistatic effects is diluting true genetic signal.

The prevalence of a few chromosome arms interacting with many other arms is of particular interest, due to the potential for one site to influence the expression of so many other sites. Jiang et al. (2017) observed a large proportion of the epistatic interactions affecting GY involved chromosomes 4A and 7D in a large population of hybrid wheat. While we did not detect a large number of interactions involving 4A, 7D was particularly important for TW. However, the interactions that they detected appear to be on the short arm of chromosome 7D, instead of the long arm as we observed. It may be that the signal detected for arms influencing multiple loci is due to the presence of functional and non-functional alleles at important upstream regulators, such as transcription factors. In this case, a non-functional transcription factor would cause the suppression of differential additive alleles. However, it appears that the loci detected to interact with many other loci in this study are not the same as those of Jiang et al. (2017).

The detection of chromosome arm interactions not identified in the homeologous marker sets suggests that single marker sets may miss important interactions. It is unclear if these interactions would have been detected if we had tested all pairwise epistatic interactions between markers. While all possible tests can be conducted, this increases the multiple testing problem drastically and may result in the loss of ability to detect even the largest effect interactions. It is unclear how large the effect sizes of a single pair of interacting loci would need to be to show up in a variance component estimated from multiple loci. While this method may not work well for a single large effect interaction, it may work well for many small effect interactions as might be expected for homeologous interactions.

It should be noted that epistatic relationships formed from the Hadamard product of covariance matrices have the property of shrinking distant relationships while emphasizing close ones. For example, two lines with an additive covariance of 0.1 will have an epistatic covariance of 0.01, whereas two lines with an additive covariance of 0.9 will have an epistatic covariance of 0.81. It may be that there are several levels of relatedness that must be considered to properly account for genetic relatedness. The pedigree is an example of a covariance estimation procedure that emphasizes close relationships and deemphasizes more distant ones. Considering both pedigree and marker based covariance matrices has been shown to be more predictive than using either alone (De Los Campos et al. 2009; Crossa et al. 2010). Other multi-kernel methods, including Reproducing Kernel Hilbert Spaces (RKHS), can be used to model these various degrees of genetic relatedness (Campos, Gianola, and Rosa 2009; Crossa et al. 2010), but may have less genetic interpretability than the method presented here.

## 6 Conclusion

The interacting pairs presented here do not have the precision to make claims of interacting genes. Nor are these interactions necessarily targets for selection. They do, however, demonstrate that there appears to be global patterns of epistasis across the genome. Seemingly additive only traits have often been shown to be under a high degree of epistasis when careful investigation is used to elucidate the trait (Carlborg et al. 2006; Forsberg et al. 2017). Some have argued that essentially all genetic variation is subject to epistasis (Huang et al. 2012; Forsberg et al. 2017), where the rest of the genome must be functional to express additive differences in alleles.

This is evident when we consider the the complexity of the cell, where no genes truly work independently of one another. In order to create the complex structure of the cell, proteins may interact with other proteins, both alike and dislike to them, to form multisubunit complexes. Therefore allelic variation alone should be sufficient to produce epistatic variation. It is merely our inability to separate this variation from “additive” variation under classic parameterizations that leads many to conclude that epistasis is not important (Hill, Goddard, and Visscher 2008; Huang et al. 2012; Huang and Mackay 2016; Forsberg et al. 2017).

Further research into this methodology might be used to identify meaningful haplotypes. Once interacting segments are identified, they can each be split into multiple pieces for further refinement of the method, while nominally increasing the number of tests performed. The low resolution epistasis mapping approach presented here emphasizes the power of using multiple genetic markers to test for interacting genomic regions, albeit at the cost of low precision.

## 7 Acknowledgments

Funding of this research was provided by the USDA National Needs Fellowship for N. Santantonio, in partial fulfillment of the requirements for a Ph.D in Plant Breeding and Genetics at Cornell University. Additionally, the field trials comprising the phenotypic data for the CNLM population were funded in part by the Hatch Project # 149-447. Genotyping was funded by the Wheat Coordinated Agricultural Project (WheatCAP). The authors thank the International Wheat Genome Sequencing Consortium for pre-publication access to IWGSC RefSeq v1.0 and Martin Mascher at the Leibniz-Institute of Plant Genetics and Crop Plant Research IPK (collaborating with IWGSC) for providing chromosome centromere positions.

## 8 Supplementary Materials

**Table S1:**
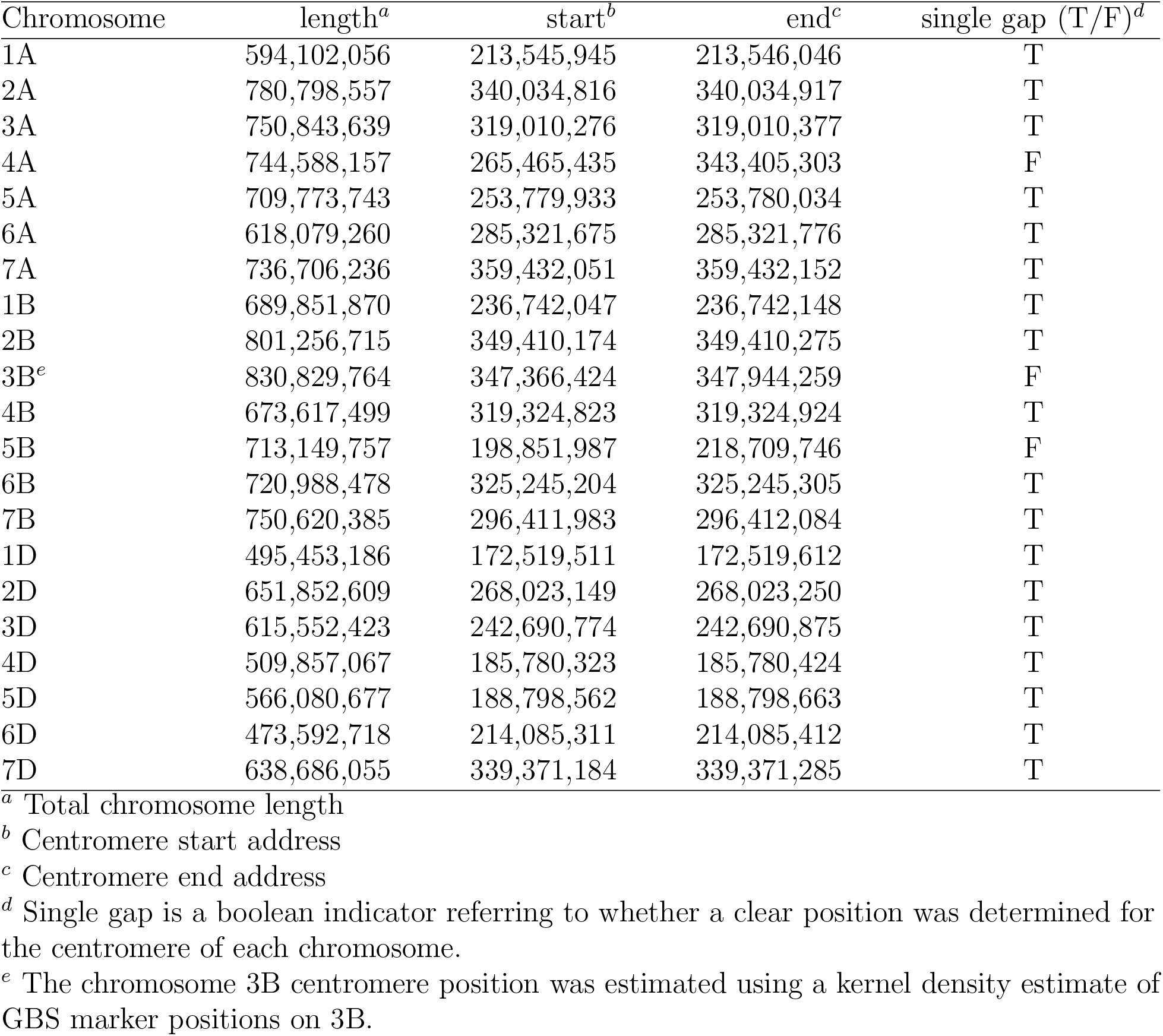
Table of centromere positions for the 21 chromosomes of hexaploid wheat based on the RefSeq v1.0 of ‘Chinese Spring’ (IWGSC 2018, accepted). These positions were provided by the IWGSC (IWGSC, personal communication, March 1, 2017)

| Chromosome | length<sup>a</sup> | start<sup>b</sup> | end<sup>c</sup> | single gap (T/F)<sup>d</sup> |
| --- | --- | --- | --- | --- |
| 1A | 594,102,056 | 213,545,945 | 213,546,046 | T |
| 2A | 780,798,557 | 340,034,816 | 340,034,917 | T |
| 3A | 750,843,639 | 319,010,276 | 319,010,377 | T |
| 4A | 744,588,157 | 265,465,435 | 343,405,303 | F |
| 5A | 709,773,743 | 253,779,933 | 253,780,034 | T |
| 6A | 618,079,260 | 285,321,675 | 285,321,776 | T |
| 7A | 736,706,236 | 359,432,051 | 359,432,152 | T |
| 1B | 689,851,870 | 236,742,047 | 236,742,148 | T |
| 2B | 801,256,715 | 349,410,174 | 349,410,275 | T |
| 3B<sup>e</sup> | 830,829,764 | 347,366,424 | 347,944,259 | F |
| 4B | 673,617,499 | 319,324,823 | 319,324,924 | T |
| 5B | 713,149,757 | 198,851,987 | 218,709,746 | F |
| 6B | 720,988,478 | 325,245,204 | 325,245,305 | T |
| 7B | 750,620,385 | 296,411,983 | 296,412,084 | T |
| 1D | 495,453,186 | 172,519,511 | 172,519,612 | T |
| 2D | 651,852,609 | 268,023,149 | 268,023,250 | T |
| 3D | 615,552,423 | 242,690,774 | 242,690,875 | T |
| 4D | 509,857,067 | 185,780,323 | 185,780,424 | T |
| 5D | 566,080,677 | 188,798,562 | 188,798,663 | T |
| 6D | 473,592,718 | 214,085,311 | 214,085,412 | T |
| 7D | 638,686,055 | 339,371,184 | 339,371,285 | T |

<sup>a</sup> Total chromosome length
<sup>b</sup> Centromere start address
<sup>c</sup> Centromere end address
<sup>d</sup> Single gap is a boolean indicator referring to whether a clear position was determined for the centromere of each chromosome.
<sup>e</sup> The chromosome 3B centromere position was estimated using a kernel density estimate of GBS marker positions on 3B.

**Figure S1:**
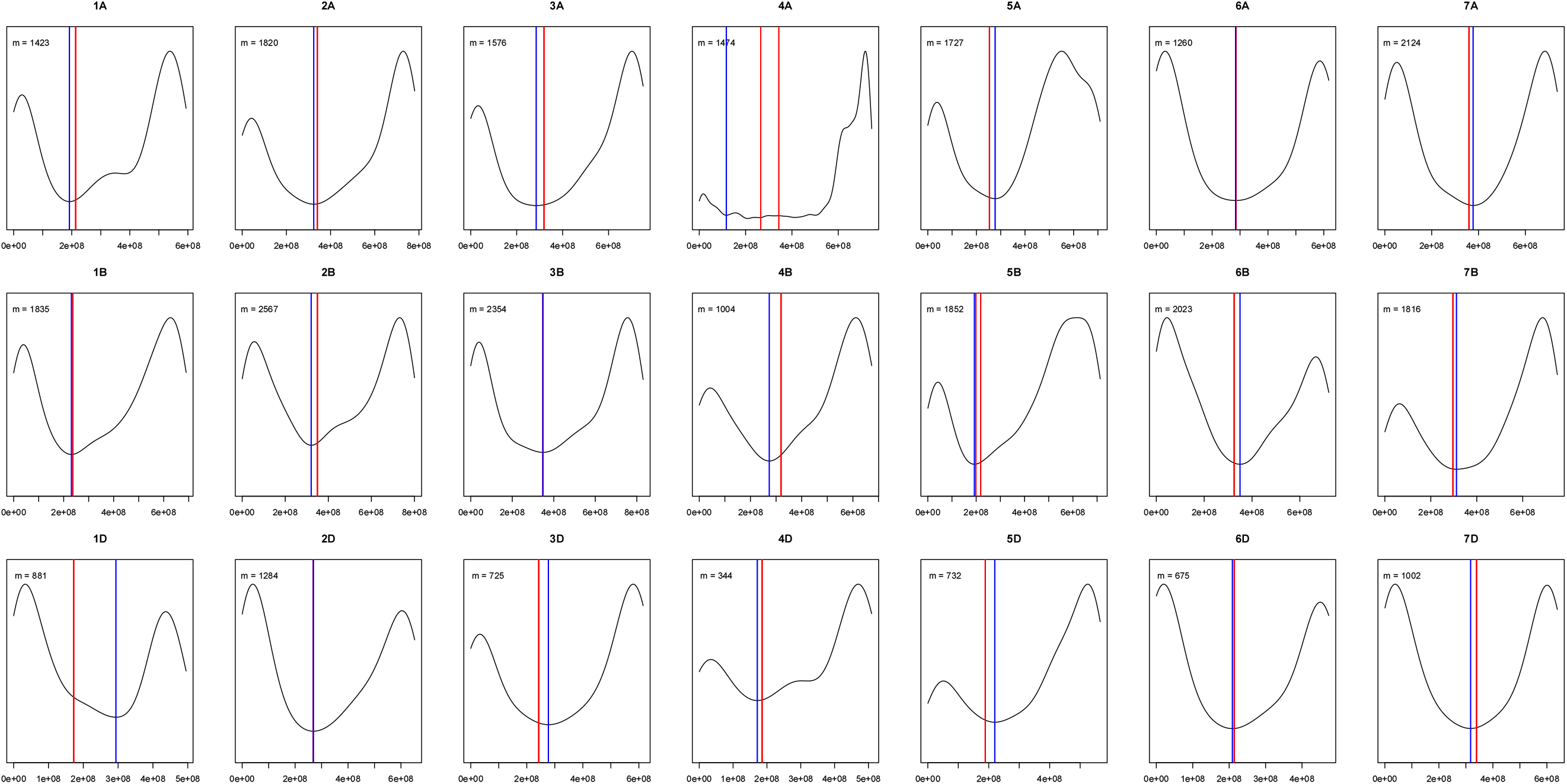
Kernel density estimation of GBS marker distribution across the 21 chromosomes of wheat. Red lines indicate the centromere interval provided by IWGSC (personal communication, March 1, 2017), and blue line indicate the centromere interval estimate based on the first derivative of the density estimate.

**Figure S2:**
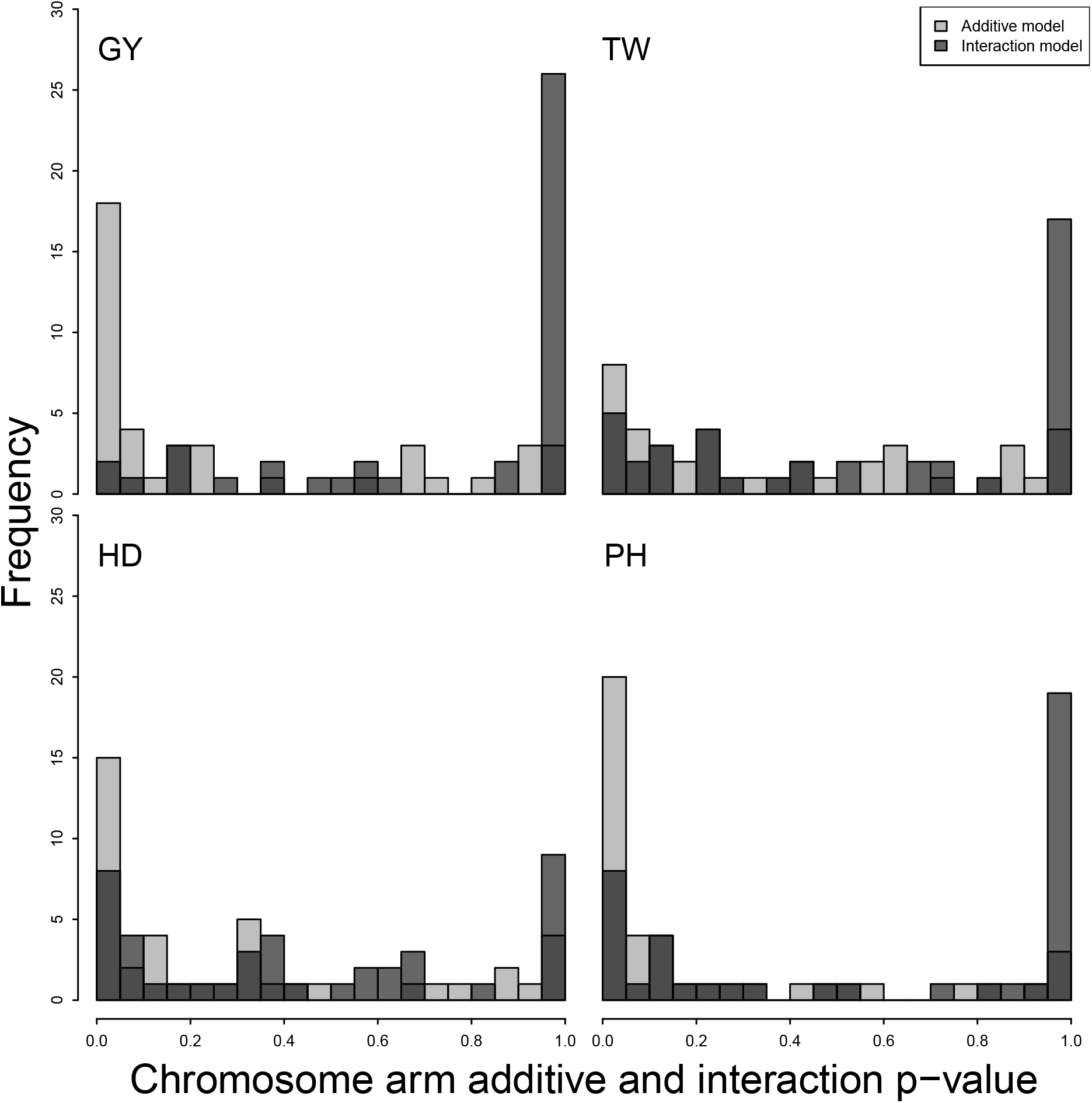
Distribution of p-values for 42 homeologous chromosome arm pair models for four traits, GY, TW, PH and HD. The p-value from the likelihood ratio test for the additive chromosome arm model is plotted in light gray, whereas the p-value from the the interaction model test is shown in dark gray.

**Figure S3:**
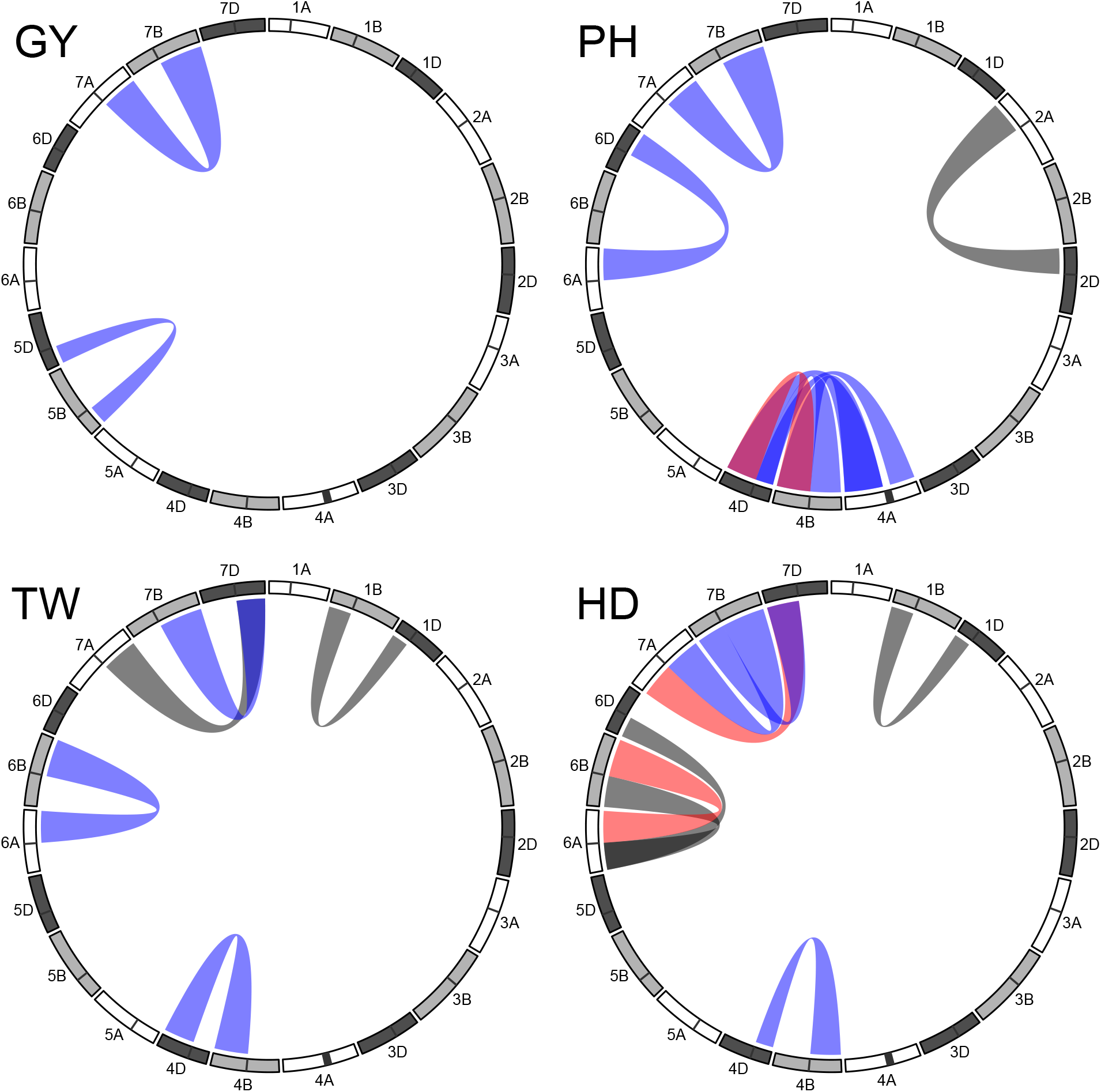
Homeologous chromosome arm interactions significant at p < 0.05. Blue and red bridges indicate interactions with a significant positive or negative correlation between the product of the additive effects and their interaction effect, respectively. Black bridges indicate significant interactions that did not have a significant correlation between additive products and the interaction effect.

**Figure S4:**
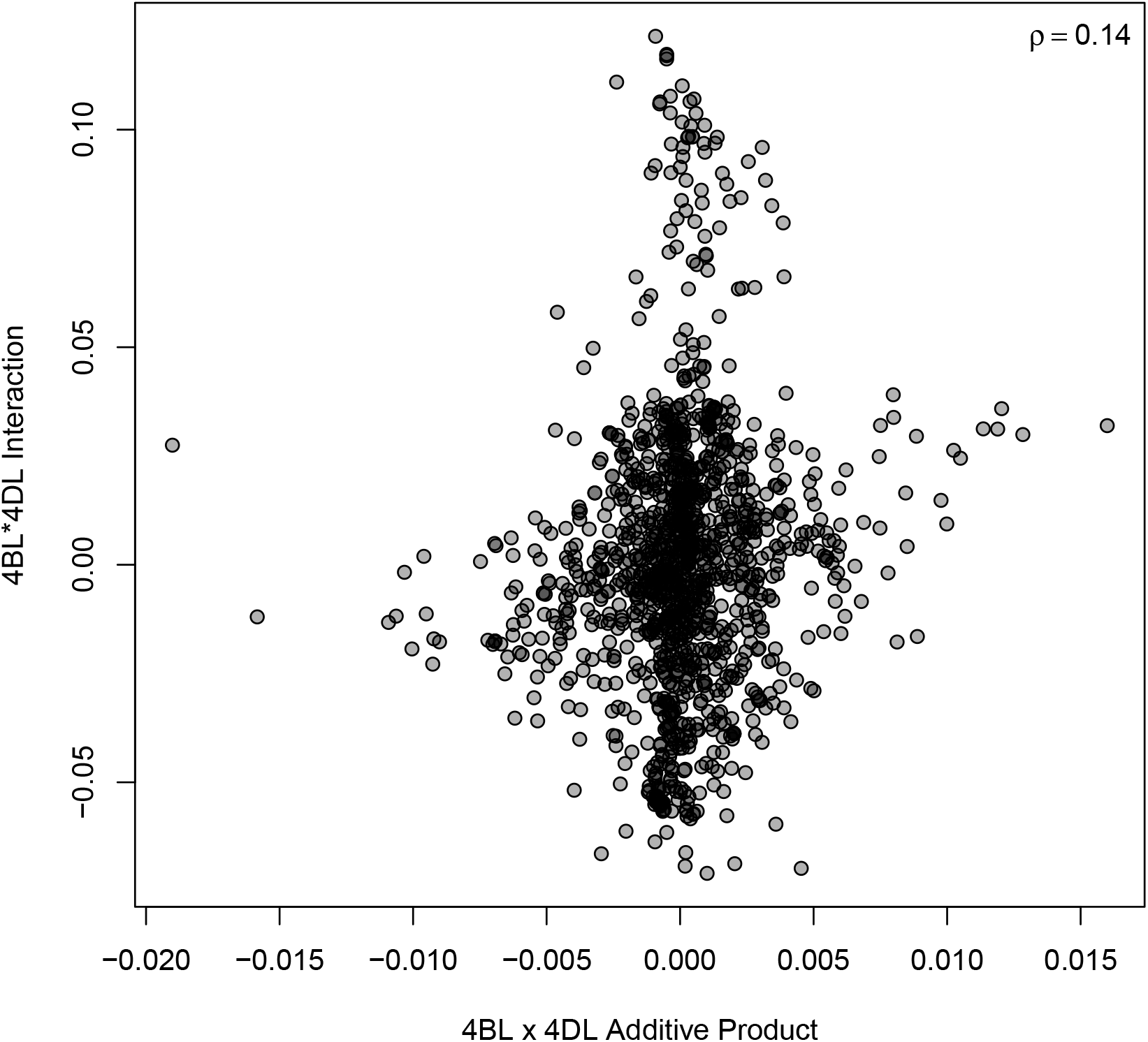
Interaction effect of chromosome 4BL by 4DL plotted against the product of the additive effects for 4BL and 4DL for TW. *ρ* indicates the Pearson correlation coefficient.

**Figure S5:**
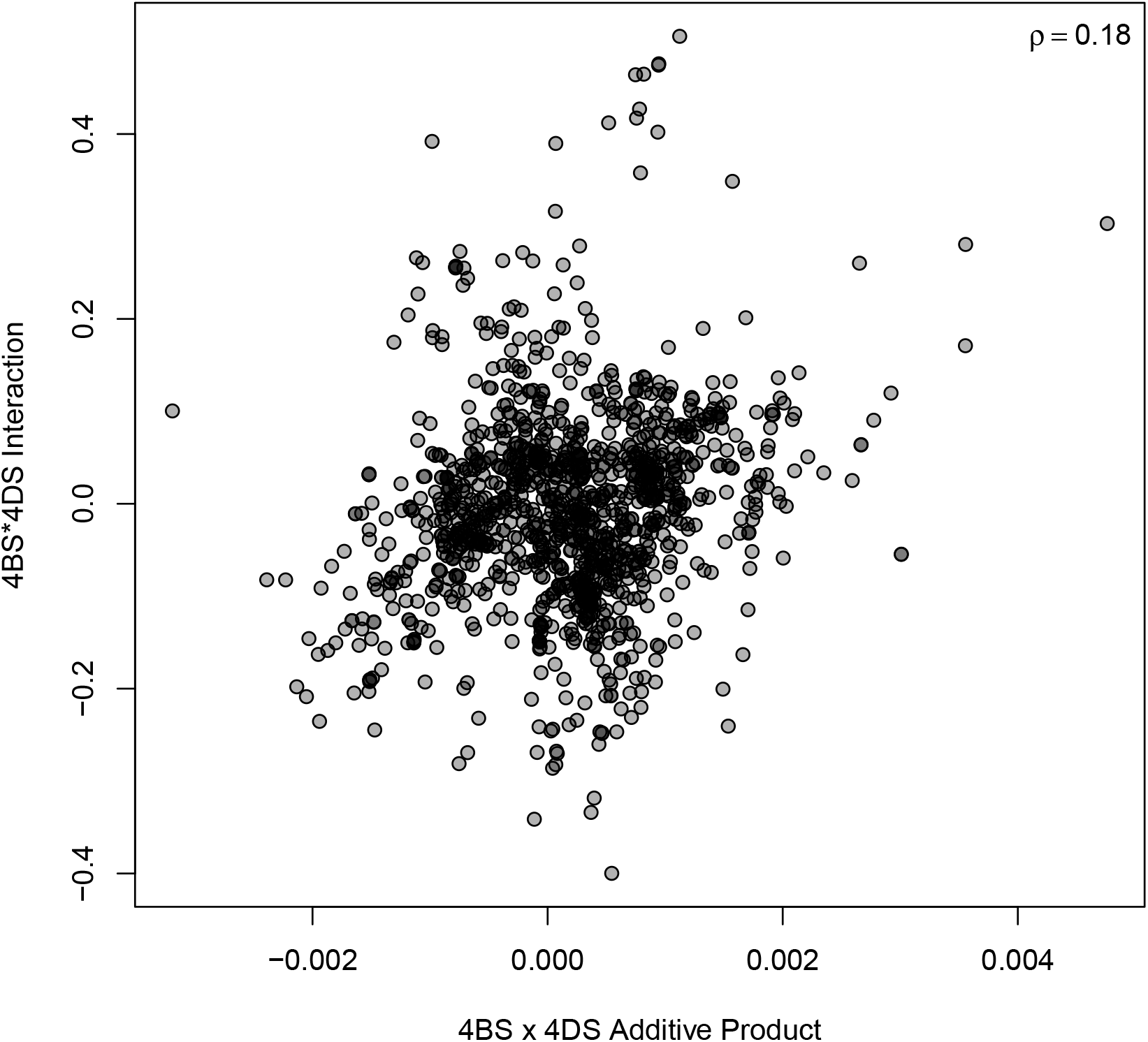
Interaction effect of chromosome 4BS by 4DS plotted against the product of the additive effects for 4BS and 4DS for PH. *ρ* indicates the Pearson correlation coefficient.

**Table S2:** Counts of significant homeologous chromosome arm interactions by arm and traits. Chomosome Arm GY PH TW HD Total 1AL

| Chromosome Arm | GY | PH | TW | HD | Total |
| --- | --- | --- | --- | --- | --- |
| 1AL | 0 | 9 | 2 | 0 | 11 |
| 1BL | 0 | 0 | 0 | 1 | 1 |
| 1BS | 0 | 4 | 3 | 2 | 9 |
| 1DS | 0 | 0 | 0 | 1 | 1 |
| 2AL | 0 | 5 | 2 | 0 | 7 |
| 2BS | 0 | 2 | 0 | 0 | 2 |
| 2BL | 0 | 1 | 1 | 0 | 2 |
| 2DS | 0 | 10 | 0 | 0 | 10 |
| 3AS | 0 | 6 | 0 | 0 | 6 |
| 3AL | 0 | 3 | 1 | 0 | 4 |
| 3BS | 0 | 0 | 2 | 1 | 3 |
| 3BL | 0 | 0 | 1 | 0 | 1 |
| 3DS | 0 | 1 | 0 | 2 | 3 |
| 4AL | 0 | 0 | 1 | 0 | 1 |
| 4BS | 0 | 7 | 0 | 0 | 7 |
| 4BL | 0 | 4 | 0 | 0 | 4 |
| 4DS | 0 | 11 | 0 | 4 | 15 |
| 4DL | 0 | 10 | 0 | 0 | 10 |
| 5AL | 0 | 0 | 2 | 0 | 2 |
| 5BS | 0 | 0 | 0 | 1 | 1 |
| 5DS | 0 | 1 | 0 | 0 | 1 |
| 6AS | 0 | 7 | 1 | 5 | 13 |
| 6AL | 0 | 1 | 0 | 0 | 1 |
| 6BS | 0 | 0 | 1 | 0 | 1 |
| 6DL | 0 | 5 | 0 | 1 | 6 |
| 7AS | 0 | 0 | 1 | 3 | 4 |
| 7AL | 0 | 7 | 1 | 0 | 8 |
| 7BL | 0 | 4 | 0 | 5 | 9 |
| 7DS | 0 | 0 | 1 | 0 | 1 |
| 7DL | 0 | 0 | 14 | 0 | 14 |
| Total | 0 | 98 | 34 | 26 | 158 |

